# Antiretroviral treatment, prevention of transmission, and modeling the HIV epidemic: why the ART efficacy and effectiveness parameter matters

**DOI:** 10.1101/228940

**Authors:** Reuben Granich, Somya Gupta, Matt Wollmers, Brian Williams

## Abstract

**Introduction:** HIV remains a major public health threat with over 75 million deaths, 2 million annual infections and over 1 million HIV-associated TB cases a year. Population-based studies suggest a marked decline in incidence, prevalence and deaths, mostly likely due to treatment expansion, in countries in East and Southern Africa. This calls into question the ART efficacy, effectiveness and coverage parameters used by many modelers to project HIV incidence and prevalence.

**Methods:** For 2015 and 2016 we reviewed global and national mathematical modeling studies regarding ART impact (with or without other HIV prevention interventions) and/or 90-90-90 on either new HIV infections or investment or both. We reviewed these HIV epidemiologic and costing models for their structure and parameterization around ART; we directly compared two models to illustrate differences in outcome.

**Results:** The nine models published in 2015 or 2016 included parameters for ART effectiveness ranging from 20% to 86% for ART effectiveness. Model 1 limits eligibility for ART initiation to 80% coverage of people living with HIV and with a CD4+ cell count below 350 cells/μL, 70% retention, and ART reduces transmission by 80%, with a derived ART effectiveness of 20%. Model 2 assumes 90-90-90 by 2020 (i.e., 73% viral suppression of estimated PLHIV), ART reduces transmission by 96% in those on ART and virally suppressed, and by 88% in those on ART but not virally suppressed with a derived effectiveness of 86% and consequent decline towards ending AIDS and HIV elimination. ART parameter selection and assumptions dominate and low ART effectiveness translates into lower impact.

**Discussion:** Using more realistic parameters for ART effectiveness suggests that through expanding access and supporting sustainable viral suppression it will be possible to significantly reduce transmission and eliminate HIV in many settings.

## Introduction

Despite progress, HIV is still a major public health threat with over 75 million deaths, 2 million infections a year and over 1 million HIV-associated TB cases a year^1, 2^. The impact of the HIV epidemic prompted an unprecedented response and we now know more about HIV than any other pathogen in history. The discovery of effective antiretroviral treatment (ART) in 1996 and subsequent evidence regarding the prevention of illness, death and transmission transformed the epidemic from an unending, unmitigated disaster into something that could be prevented and even ended someday^3-9,10^. The potential for expanded access to ART to curb the epidemic and the concept of treatment as prevention was introduced in 2006^5^ and later formalized by the World Health Organization (WHO)^11^. Over the past decade, the global and local HIV strategy has shifted from “test-and-wait” to “test-and-treat” focusing on achieving the 90-90-90 target (73% of people on ART and virally suppressed) and ending AIDS (defined as universal treatment with less than one AIDS case and one AIDS related death per 1000 population) ^12 13, 14^.

Policy discussions around treatment-as-prevention have focused on how to increase access to testing and how early to provide diagnosis and treatment. Global HIV leaders and stakeholders turned to modeling to answer key questions about the risks and benefits of expanding access to treatment. Before and after the prevention impact of ART was understood, researchers used models to explore the impact of possible ART expansion scenarios^15-19^. While a few models explored expansion of ART access beyond existing WHO guidelines, most models limited testing and treatment to those who were severely immunocompromised^19^. Similarly, traditional costing efforts could be classified as “doomsday costing” insofar as they took a health sector perspective and only looked at the costs of providing earlier treatment while ignoring the potential prevention benefits and cost savings of earlier treatment20, 21. More modern “second generation” approaches to economic modeling took into account the prevention impact of scaling up treatment along with other interventions^18,22-24^. These models explored treatment as prevention of illness, death, and transmission. In some cases, the models were combined with a costing framework to examine the costs, cost benefits, and costs savings of various scale-up scenarios^22-25^. The dominant model (GOALS) used by the Joint United Nations Programme on HIV/AIDS (UNAIDS), the Global Fund and the United States government and many countries now includes the prevention impact of ART and is used to determine the health and transmission impact, costs and cost-savings for various HIV response scenarios^23^. The UNAIDS estimates uses GOALS that incorporates data from available surveys and other surveillance information and makes forward projections of incidence, prevalence and resources needs according to their financial framework categories^23^. In the 2016 the UNAIDS HIV Update and in the UNAIDS resource needs projections, the incidence and prevalence were reported as being stable for the 5 years from 2010 to 2015 in all regions of the world except for Eastern Europe where the rates were increasing1, 23. During this time the world spent an estimated US$200 Bn on attempting to control HIV and the conclusion that can be drawn from the report is that the significant investment has had little or no impact on incidence or mortality. If true, this has major implications for future resource needs as well as for global HIV control strategy since the impact of treatment appears to be far less than expected. However, there are reasons to question the flat-line UNAIDS estimates of incidence and prevalence as these results contrast with other models and the scientific evidence regarding the potential impact of ART and other prevention interventions. Recent population-based studies from a number of countries suggest a marked decline in incidence, prevalence and deaths, mostly likely due to treatment expansion, in many countries in East and Southern Africa^26^.

## Methods

The marked contrast in model outcomes, one predicting flat-line incidence and prevalence in four of five regions^1, 23^ while others projecting a steady decline to elimination, prompted us to explore the importance of ART efficacy, effectiveness and coverage parameters. Specifically, for 2015 and 2016, we reviewed global and national mathematical modeling studies that looked at impact of ART (with or without other HIV prevention interventions) or 90-90-90 on either new HIV infections or investment or both. We reviewed these HIV epidemiologic and costing models for their structure and parameterization around ART.

## Results

Table 1 describes the 9 models, including the available parameters used to derive ART effectiveness expressed in terms of percentage reduction in HIV transmission. The modeling parameters for ART effectiveness by 2020 ranged from 20% to 86% for ART effectiveness. This disparity in ART Effectiveness is further highlighted in Figure 1 that shows the comparison between the GOALS and SACEMA models for Mozambique (other model comparisons not shown). The GOALS model forms the basis for UNAIDS estimates and gives a more pessimistic prediction of the trend in incidence and mortality when compared with the SACEMA model in the left panel of the figure. The GOALS model limits eligibility for ART initiation and assumes that at full coverage 80% of those infected with HIV and with a CD4+ cell count below 350 cells/μL. With these assumptions about half of all those infected with HIV will be on ART which, when coupled with the lower estimate of transmission impact, yields a derived ART effectiveness of only 20%. The SACEMA model, on the other hand, assumes that we reach 90% ART coverage by 2020 and then continue to roll-out ART at the same rate while ART reduces transmission by 96%; this results in a derived ART effectiveness of 86% and a much more optimistic forecast. The underlying models are similar. To match the GOALS model the SACEMA model would need to adjust the parameters downward to assume that only 65% of those infected with HIV are on ART at full coverage instead of 90% and ART reduces transmission by 65% instead of 96%. Clearly, these two parameters are the critical determinants of the impact of treatment on incidence and mortality. Comparisons with other models show similar results with the ART effectiveness parameter driving outcomes.

**Table 1:** Description of Models with Calculated ART Effectiveness.

| STUDY | SETTING | PARAMETERS | ART EFFECTIVENESS |
| --- | --- | --- | --- |
| **1. Williams (SACEMA model)**, Current HIV AIDS Research 2015<br>Epidemiological Trends for HIV in Southern Africa: Implications for Reaching the Elimination Targets | Southern Africa | Proportion on ART virally suppressed: 90%<br>Reduction in transmission on ART<br>* virally suppressed 96%<br>* not virally suppressed 88%<br>Full coverage (2020): People at risk tested on average twice a year and started on treatment immediately | **Effectiveness: 86%**<br>**Calculations (for 2020):**<br>On ART: 90%<br>Not on ART: 10%<br>Of those on ART percentage virally suppressed: 90%<br>Of those on ART percentage not virally suppressed: 10%<br>On ART and virally suppressed transmission reduced: 96%<br>On ART and not virally suppressed transmission reduced: 88%<br>Effectiveness:<br>$1-(0.9*(0.9*0.04+0.1*0.12)+0.1)$<br>=0.86 |
| **2. Smith (Imperial model)**, Lancet HIV 2016<br>Maximizing HIV prevention by balancing the opportunities of today with the promises of tomorrow: a modeling study | South Africa | Efficacy (protection afforded by perfect use of a product) of early ART: **85%**<br>Effective coverage (proportion of people who fully adhere to a product such that they benefit from its protection): Constant: 0%; medium: 40%; maximum: **60%** | **Effectiveness: 51%**<br>**Calculations (for 2020):**<br>On ART: 60%<br>Not on ART: 40%<br>On ART transmission reduced: 85%<br>Effectiveness:<br>$1-(0.6*0.15+0.4) = 0.51$ |
| **3. McGillen (Imperial model)**, Lancet HIV 2016<br>Optimum resource allocation to reduce HIV incidence across sub-Saharan Africa: a mathematical modelling study | 18 countries from sub-Saharan Africa (80% of adult HIV burden in the region) | Effectiveness of early ART as prevention (reduction in risk of onward transmission): **70%**<br>Note: Early ART refers to a prevention method comprising outreach testing programmes and the offer of treatment to all PLHIV.<br>Achievable coverage: 33% among heterosexual men and low-risk women and 66% among MSM and FSW (this is the coverage of early ART for PLHIV who have not already presented for care i.e. their CD4 is above 200 initially and above 350 later when ART eligibility shifted)<br>Proportion of people living with HIV who are virally suppressed: **63%** | **Effectiveness: 44%**<br>**Calculations (for 2020):**<br>On ART: NA<br>Not on ART: NA<br>On ART virally suppressed: 63%<br>On ART not virally suppressed: NA<br>On ART transmission reduced: 70%<br>Effectiveness:<br>$1-(0.63 \times 0.3 + 0.37) = 0.44$ |
| **4. Stover (GOALS model)**, PLoS ONE 2016<br>What Is Required to End the AIDS Epidemic as a Public Health Threat by 2030? The Cost and Impact of the Fast-Track Approach | 45 countries (86% of new infections globally) | 95% reduction in infectiousness among those virally suppressed<br>Adult ART 2020 coverage: 81% (90% started, 90% retained); 90% of them are retained and 90% are virally suppressed<br>Adult ART 2030 coverage: 90% (95% started, 95% retained); 95% of them are virally suppressed<br>Eligibility for treatment expands to all PLHIV by 2018 | **Effectiveness: 62%**<br>**Calculation (for 2020):**<br>On ART: 81%<br>Not on ART: 19%<br>Of those on ART percentage virally suppressed: 81%<br>Of those on ART percentage not virally suppressed: 19%<br>On ART and virally suppressed transmission reduced: 95%<br>On ART not virally suppressed transmission reduced: NA<br>Effectiveness:<br>$1-(0.81 \times (0.81 \times 0.05 + 0.19) + 0.19) = 0.62$ |
| **5. Kripke (DMPPT 2.1 model)**, PLoS ONE 2016<br>Impact and Cost of Scaling Up Voluntary Medical Male Circumcision for HIV Prevention in the Context of the New 90-90-90 HIV Treatment Targets | Lesotho, Malawi, South Africa, Uganda | “ART effect ” parameter (ratio of infectiousness with ART to without ART) was used to model the level of viral suppression: base value was 0.25 (till 2015) and reduced to 0.1 (by 2020) and 0.05 (by 2030) under 90-90-90 scenario<br>Adult ART 2020 coverage: 81%<br>Eligibility for treatment expands to all PLHIV by 2017 | **Effectiveness: 73%**<br>**Calculations (for 2020):**<br>On ART: 81%<br>Not on ART: 19%<br>On ART transmission reduced: 90%<br>Effectiveness:<br>$1-(0.81 \times 0.1 + 0.19) = 0.73$ |
| **6. Korenromp (GOALS model)**, PLoS ONE 2015<br>Impact and Cost of the HIV/AIDS National Strategic Plan for Mozambique, 2015-2019—Projections with the Spectrum/Goals Model | Mozambique | ART reduces infectivity of PLHIV by **80%**, as an average effectiveness between recent studies including a 96% reduced infectivity found in a clinical trial across multiple—mainly developed, western—countries with very high adherence [Cohen NEJM 2011; Attia AIDS 2009], a 38% reduction in a high-coverage ART program in rural South Africa [Tanser Science 2013]; 85% suppression observed in Swaziland [SHIMS 2010-12; Justman CROI 2013]<br>Scenario ‘current targets’: ART is scaled-up from 56% to 76% of adults with CD4 <350 in North region, from 65% to 81% in Center, and from 57% to 85% in South; additionally eligibility includes TB/HIV-co-infected adults and pregnant women (from 2012 and 2014, respectively), in all scenarios irrespective of CD4 count.<br>Scenario ‘Accelerated scale-up’: ART is further scaled-up to 85% of eligible PLHIV with CD4 <350 and all FSW irrespective of CD4 count<br>Retention on ART, at 3 years after enrolment: 52% in current targets scenario and 70% in Accelerated scale-up scenario | **Effectiveness: 20%**<br>**Calculations (for 2020):**<br>On ART: 35% (imputed)<br>[85% of those <350 CD4 cell count (i.e. 25% of PLHIV), ART for all pregnant women, female sex workers, TB/HIV]<br>Not on ART: 65%<br>Retention on ART: 70%<br>On ART transmission reduced: 80%<br>Effectiveness:<br>$1-((0.35 \times 0.7 \times 0.2) + (1 - 0.35 \times 0.7)) = 0.20$ |
| **7. Walensky (CEPAC model),** Ann Intern Med 2016<br>The Anticipated Clinical and Economic Impact of 90-90-90 in South Africa | South Africa | UNAIDS Target strategy: 73% suppression in 5 years from 24% current<br>HIV transmission rates by disease stage and viral load: 0.16–9.03/100 person years<br>Mean ART efficacy, % virologic suppression at 48 weeks: 72% | **Effectiveness: 52%**<br>**Calculations (for 2020):**<br>On ART: 81%<br>Not on ART: 19%<br>Of those on ART percentage virally suppressed: 90%<br>Of those on ART percentage not virally suppressed: 10%<br>On ART and virally suppressed transmission reduced: 72%<br>On ART not virally suppressed transmission reduced: NA<br>Effectiveness:<br>$1-(0.81*(0.9*0.28+0.1*1)+0.19) = 0.52$ |
| **8. Olney (Imperial model),**<br>Lancet HIV 2016<br>Evaluating strategies to improve HIV care outcomes in Kenya: a modelling study | Kenya | • Infectiousness of HIV-positive, on ART and virally suppressed: 0.1 (estimate)<br>• Proportion of individuals initiating ART who adhere to ART and achieve viral suppression: 86% | **Effectiveness: 63%**<br>**Calculations (for 2020):**<br>On ART: 81% (assumption based on UNAIDS 90-90-90 target)<br>Not on ART: 19%<br>Of those on ART percentage virally suppressed: 86%<br>Of those on ART percentage not virally suppressed: 14%<br>On ART and virally suppressed transmission reduced: 90%<br>On ART not virally suppressed transmission reduced: NA<br>Effectiveness:<br>$1-(0.81*(0.86*0.1+0.14)+0.19)= 0.63$ |
| **9. Hontelez (STDSIM model),** AIDS 2016<br>Changing HIV treatment eligibility under health system constraints in sub-Saharan Africa: investment needs, population health gains, and cost-effectiveness | 10 sub-Saharan African countries (80% regional burden) | • ART reduces infectiousness of HIV by 90%<br>• Under 90-90-90 scenario, 81% ART coverage and 73% viral suppression among people living with HIV achieved | **Effectiveness: 73%**<br>**Calculations (for 2020):**<br>On ART: 81%<br>Not on ART: 19%<br>On ART transmission reduced: 90%<br>Effectiveness:<br>$1-(0.81*0.1+0.19)= 0.73$ |

\*PLHIV –people living with HIV, MSM –men who have sex with men, FSW –female sex workers, NA – not available
Note: When data are not available, we assume 0% reduction in transmission

**Figure 1:**
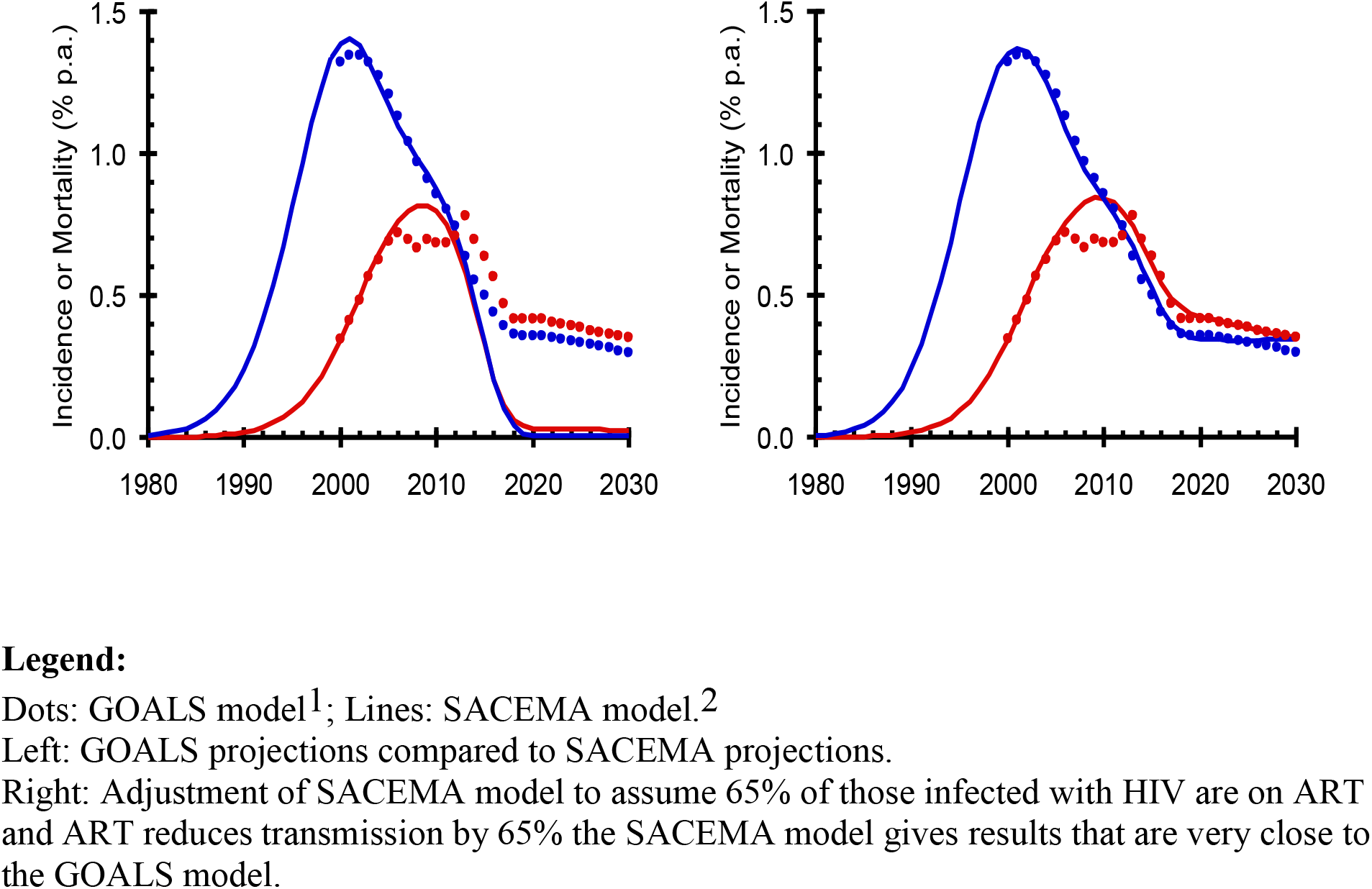
Annual HIV incidence (blue) and mortality (red) for Mozambique.

## Discussion

Modeling plays an important role in exploring potential outcomes while illustrating what data are missing for more accurate predictions. While parameterization relies on studies and other surrogate information, it is critical to be clear about the judgments involved in selecting critical values for interventions such as ART coverage and effectiveness. Studies now show that over 90% of people will agree to an HIV test if it is offered, that people started on ART adhere to a very high level, that ART suppresses the virus and reduces transmission by close to 100%, and that retention is probably much higher than predicted. Additionally, ART costs have fallen significantly and high levels of ART and viral suppression can be achieved. Being clear about what parameters to use and how they are applied to the epidemiology and costing is critical when using models to guide the HIV response. Our brief review suggests that many of the current models have likely underestimated the impact of ART. This overly conservative assessment could account for past decisions to under invest in expanding access to treatment while using remaining resources for other budget categories^27^. Using more realistic parameters around ART suggests that as we expand access and support the achievement of sustainable viral suppression it will be possible to significantly reduce transmission and eliminate HIV in many settings.

## Footnotes

1. Korenromp, E., et al., Impact and Cost of the HIV/AIDS National Strategic Plan for Mozambique, 2015-2019—Projections with the Spectrum/Goals Model. PLOS One, 2016. 10 (11): e0142908. doi:10.1371.

2 Williams, B.G., et al., Epidemiological Trends for HIV in Southern Africa: Implications for Reaching the Elimination Targets. Current HIV/AIDS reports, 2015. 12: p. 1-11.

